# Single cell RNA-seq reveals protracted germ line X chromosome reactivation dynamics directed by a PRC2 dependent mechanism

**DOI:** 10.1101/2023.11.06.565813

**Authors:** Yaqiong Liu, Xianzhong Lau, Munusamy Prabhakaran, Carlos M Abascal Sherwell Sanchez, Daniel Snell, Mahesh Sangrithi

## Abstract

Initiating soon after PGC specification, female germ cells undergo reactivation of the silenced X chromosome during genome wide reprogramming. However, the kinetics and dynamics of XCR *in vivo* have remained poorly understood. To address this here we perform a global appraisal of XCR using high-dimensional techniques. Using *F*_*1*_ *B6 v CAST* mouse embryos, we perform a detailed assessment, applying single-cell RNA-seq and chromatin profiling on germ cells purified from E10.5 to E16.5. While scRNA-seq profile showed that male and female germ cells are transcriptionally indistinct at E11.5, they are sexually dimorphic by E12.5, diverging further through development to E16.5. With allelic resolution, we show that the reactivating X chromosome is only partly active at E10.5, then reactivates gradually and reaches near parity in output to the constitutively active X chromosome at ∼E16.5 when developing oogonia are meiosis prophase I. Crucially, we show that sexually dimorphic dosage compensation patterns observed in germ cells, occur in tandem with an increase in the allelic proportion from the reactivating X chromosome. While *Xist* is extinguished from E10.5, the epigenetic memory of earlier XCI in female cells persists much longer, likely from self-sustained PRC2 complex (*Ezh2 / Eed / Suz12*) function. The reactivating X chromosome is enriched in the epigenetic silencing mark H3K27me3 at E13.5, which is removed by E16.5 permitting gene expression. Our findings link XCR, along with functional regulation of PRC2 in promoting female meiosis.

## Introduction

The accurate transmission of genomic information over generations involves the complex regulation of chromatin in germ cells. This process, germ cell genome wide reprogramming (GWR), initiates soon after primordial germ cell (PGC) specification very early in mammalian embryonic development. In the mouse, PGCs arise from the proximal epiblast at ∼E6.25 - 6.75. By around E7.25, around 30-40 PGCs are present in the developing embryo, and continue to divide mitotically increasing in numbers. From E8.5 – 9.5 early PGCs migrate along the hindgut endoderm and reach the developing gonadal primordia by ∼E10.5. At approximately E11.5 gonadal sex determination occurs, where expression of the Y-chromosome encoded *Sry* transcription factor in the gonadal pre-Sertoli cells initiates male gonadal fate and formation of a testis (Koopman et al., 1990). Conversely in females, the absence of the Y chromosome instigates the formation of an ovary. Thereafter gonadal PGCs assume sex-specific developmental fates.

GWR initiates soon after PGC specification at E6.75 and is sustained cell autonomously. It involves re-expression of the pluripotency network of genes (including *Pou5f1* (aka. *Oct4*), *Sox2, Nanog, Prdm14*), and repression of the somatic differentiation program, global DNA demethylation and removal of parental imprints (Tang et al., 2016). PGC further undergo extensive chromatin changes and remodelling of histone marks. Specifically female PGCs start to reverse X chromosome inactivation (XCI) established earlier at E5.5, i.e. X-chromosome reactivation (XCR).

The timing and dynamics of germ cell XCR are not fully understood. Early studies surmised that reactivation of X-linked genes in PGCs mostly occurred after they reached the genital ridge (∼E11.5) (de Napoles et al., 2007; Johnston, 1981; Kratzer and Chapman, 1981; Monk and McLaren, 1981; Tam et al., 1994). Landmark experiments performed in single germ cells obtained from embryos, using reverse transcription followed by polymerase chain reactivation (RT-PCR), *Xist* RNA-FISH and immunofluorescence microscopy, had indicated that gene expression from the reactivating X-chromosome initiates in nascent PGCs (Sugimoto and Abe, 2007). While limited by a small number of X-linked genes and cells being assayed, they showed that the process of XCR in PGCs commences earlier, during the pre-gonadal phase of development, progresses gradually and was incomplete at E14.5. Similar results were reported in recently in PGC-like cells (PGLCS) derived *in-vitro* from mouse embryonic stem (ES) cells, highlighting that incorrect rapid XCR kinetics had resulted in limited meiotic potential (Severino et al., 2022).

XCI in the embryo is enacted by expression of the long non-coding RNA *Xist* (Kay et al., 1993; Penny et al., 1996). *Xist* directly represses transcription by evicting the RNA polymerase transcription machinery, while also initiating a cascade of epigenetic processes including the loss of H3K4me3, recruitment of histone deacetylases to remove activating histone marks (e.g. H3K27ac), and the *Polycomb repressor complex* PRC1 / 2 complexes which deposit silencing histone marks H2AK119ub1 and H3K27me3 respectively (reviewed in Loda et al. (2022)). Later in XCI, CpG-islands (CGI) of genes subject to XCI are methylated, and the inactivated X (*Xi*) is compacted forming heterochromatin. Expression of *Xist* RNA is downregulated in female PGCs from around E7.75 - E9.5, and is extinguished by ∼E11.5 (de Napoles et al., 2007; Sangrithi et al., 2017; Sugimoto and Abe, 2007). While histone marks are globally reorganized during PGC development, the extent to which PRC 1 / 2 related repressive marks deposited as a specific consequence of XCI in the epiblast are remodelled remains poorly understood. For instance global changes to H3K27me3 levels are known to occur as part of PGC development, and these can be challenging to disentangle from more specific regulation of the mark relating to XCI (Lowe et al., 2022; Saitou et al., 2012).

It has been shown previously that sexually dimorphic dosage compensation states arise during germ cell GWR, which is dependent on the number of X-chromosomes present (Sangrithi et al., 2017). During GWR, females (XX) show X chromosome to autosome (X:A) expression ratios greater than 1, while this was below 1 in males (XY). Strikingly the elevated X:A ratios in oogonia persists from E11.5 until entry into meiosis. Hence elevated X:A ratios in females may be inherently linked to the sexually dimorphic developmental fates of germ cells, including timely meiotic-entry or / and the re-establishment of transcription on the reactivating X (Sangrithi and Turner, 2018). Indeed dosage differences in X-linked genes could promote germ cell sexual dimorphism and meiosis entry and / or progression itself in females, via the involvement of X-linked genes in these processes and possibly XCR itself.

In this article we perform a global analysis of XCR dynamics of XCR using single-cell RNA sequencing on developing mouse germ cells *in vivo* during the course of GWR. We leverage on a genetically tractable murine model that enables clear delineation of the constitutively active and reactivating X chromosomes for females. Using this model we describe a precise allele-specific map of XCR, that charts the overall kinetics of this classic epigenetic process, but also the dynamics of individual X-linked genes. This study demonstrates that XCR in female germ cells initiates before E10.5 and accelerates while a sexually dimorphic transcriptome is established. We demonstrate that transcription from the reactivating X chromosome reaches close to parity with the constitutively active X chromosome only at ∼E16.5 when developing oogonia are in zygonema / early pachynema. *Xist-*dependent silencing that is present in early PGCs, then gives way to persisting repression of X-linked genes due to histone 3 lysine-27 trimethylation (H3K27me3) later in gonadal germ cells where direct *Xist* activity has ceased. H3K27me3 marks remain at the transcription-start sites (TSSs) and gene-bodies of repressed X genes on the reactivating X chromosome at meiotic entry (pre-Leptonema) in E13.5 females germ cells, with germ line genes expressed thereafter showing a dynamic reduction in H3K27me3 at E16.5. We posit XCR as a mechanism of meiotic upregulation in females.

## Results

### Transcriptional divergence of mouse germ cells occurs after E12.5

We developed a model to examine germline XCR, by combining a classical *F*_*1*_ genetics and single-cell RNA-sequencing (scRNA-seq). Leveraging on an interspecific cross of *Mus castaneus* (*CAST*) males with reference strain C57B6J (*B6*) female mice, we performed single-cell RNA-seq on *F*_*1*_ female (X_*CAST*_X_*B6*_) and male (X_*B6*_Y) embryonic germ cells from E10.5 to E16.5 stages. The *CAST* strain is highly polymorphic in relation to the reference *C57B6NJ* strain, on average containing up to 1 single-nucleotide polymorphism (SNP) for every 150bp in the genome (Keane et al., 2011). We further introduced an *Xist*-null (*XistΔ*) allele into our experimental strategy to skew X-inactivation toward the *CAST* X chromosome (Marahrens et al., 1998). By crossing *Xist*_*+/Δ*_ C57B6J (*B6*) females (carrying *Oct4-EGFP* transgene) and *CAST* males, we derived *F*_*1*_ female (X_*CAST*_X_*B6-XistΔ*_) and male (X_*B6*_Y) embryos, from which embryonic gonads were obtained daily from E10.5 to E16.5 (Yeom et al., 1996; Yoshimizu et al., 1999). Single *EGFP* positive germ cells were sorted using fluorescence-activated cell sorting (FACS) and processed to generate high-quality scRNA-seq libraries using the SMART-seq2 protocol (Picelli et al., 2014). To distinguish allele-specific expression in *F*_*1*_ embryos, reads were first aligned to a reference genome with SNPs N-masked to minimize bias arising during alignment (Degner et al., 2009). Reads were then further assigned specifically to each of the parental (*B6* and *CAST*) genomes respectively to obtain allele-specific counts (**Figure 1A**; please refer to **Methods** for details) (Krueger and Andrews, 2016). Where possible we sought to include at least 20 germ cells from at least two individual embryos at each time point and sex. Following stringent quality control checks, we retain 681 single-cell transcriptomes for further analysis (**Supplementary Figure 1A**). These include 333 male and 348 female cells (**Supplementary Figure 1B**).

**Figure 1.**
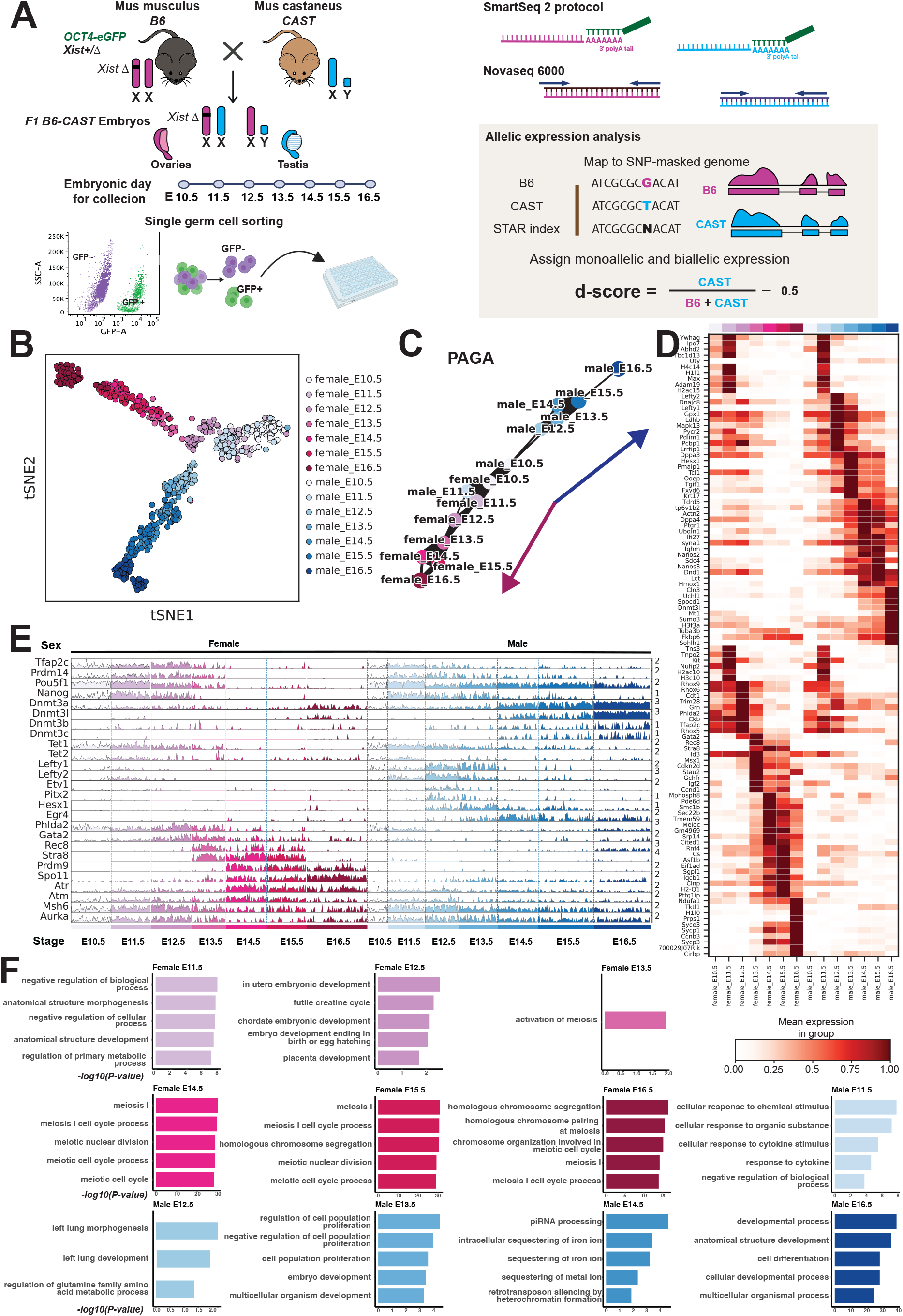
Transcriptome profiling of mouse germ cells by scRNA-seq. A. Illustration of experimental design. B. *tSNE* analysis of a total of 681 germ cell scRNA-seq data. Cells are coloured according to sex and developmental stages. The number of cells at each stage and sex are summarised in the **Supplementary Figure 1**. C. Partition-based graph abstraction (PAGA) plot representing a graph of inferred connectivity among all cell clusters. Line thickness indicates the strength of cluster connections. D. Heatmap of top 10 DE genes at different embryonic stages in female and male germ cells. E. A tracks-based bar-plot depicting the expression levels of key genes involved in PGCs, DNA methylation regulation, prospermatogonial / oogonial development, and meiosis in females and males. F. Top GO:BP (Gene Ontology for Biological Process) terms with *p*-values for different embryonic stage germ cells.

First, we performed dimensionality reduction on the dataset. The scRNA-seq profiles reveals that male and female germ cells appear transcriptionally indistinct prior to E12.5 (**Figure 1B**). Following gonadal sex determination at E11.5, sexually dimorphic transcriptomes are apparent by E12.5, which diverge further with continued development to E16.5. We sexed our samples based on the expression of Y-chromosome encoded genes in the data which is typically only detected males, and concordant with PCR based genotyping that was performed at the time of sample collection (**Supplementary Figure 1C**). We utilized partition graph abstraction (PAGA) analysis to visualize connectivity and relatedness between cell groups in an unbiased manner. The PAGA graph emphasizes the divergence of the male and female germ line after E11.5, further lending weight that our data reliably captures the establishment of sexually dimorphic transcriptomes in germ cells (**Figure 1C**).

We next performed rigorous differential expression (DE) testing to examine transcriptional changes arising during the establishment of germ cell transcriptional identity (**Figure 1D**). Expression of the *Tet* enzymes (*Tet1* and *Tet2*), that regulate DNA demethylation was observed in both male and female PGCs at E11.5, consistent with germ cells reaching a methylation base state by E13.5 (Hackett et al., 2013; Popp et al., 2010; Seisenberger et al., 2012). Male germ cells then begin to express the *Lefty* genes (*Lefty1* and *Lefty2*), *Pitx2* and the transcription factor *Etv1* that is important in prospermatogonial development. In females, we observe *Phlda2*, along with transcription factors and *Gata2*. Consistent with the pre-leptonema stage, female germ cells subsequently begin to upregulate meiotic genes at E13.5, including *Rec8, Stra8, Prdm9, Spo11* and genes with roles in meiosis I prophase thereafter (**Figure 1E** and **Supplementary Figure 1D**). Together these culminate in the establishment of pro-spermatogonia and oogonia respectively by E16.5, confirmed with gene ontology (GO) enrichment scoring (**Figure 1F**). We next turned our focus toward understanding germ cell XCR in females.

### Global X chromosome reactivation dynamics

In our experimental model, the *XistΔ* allele skews XCI invariably toward the *CAST* X chromosome. The *CAST* X-chromosome thus is always inactivated in all cells in the epiblast, and hence will also be the X chromosome that undergoes XCR in germ cells. Hence expression specific to the *CAST* X-chromosome in our *F*_*1*_ germ cells can be used to accurately chart XCR. We computed allele-level counts and calculated an allelic deviation score (‘*d* score’), as the ratio of B6 reads to the total number of informative allelic reads (i.e. *d = CAST / (B6 + CAST) - 0.5)* for each gene (Xu et al., 2017) (see **Methods**). A negative *d-score* corresponds to a bias toward expression from *B6* alleles, while positive values corresponding to a bias toward *CAST* alleles, with *d-scores* ∼0 being indicative of equal allelic expression. Instances where the *d-*scores are less than -0.4 or greater than 0.4 represent monoallelic expression bias. To test the assumptions of our experimental model, we first turned to inspecting allelic balance in E11.5 female gonadal somatic cells obtained as a control. Consistent with the *CAST* X chromosome being inactivated invariably in somatic cells, we substantiate a strong expression bias toward alleles on the *B6* X chromosome (median *d*_*ChrX*_ = -0.42) in E11.5 female somatic cells. (**Supplementary Figure 2A**). This affirms that our approach is efficient in identifying allelic expression deviation expected in XCI.

Next, we turned to studying allelic expression globally in female germ cells. We computed global *d-*scores for each chromosome in all cells in our dataset. In more detail, we see that male germ cells have *d-scores* of ∼-0.5 throughout, consistent with these cells only having a *B6* X chromosome. In females, we observe that E10.5 female PGCs had a *d-score* of -0.2 for expression from the X chromosome, with a minimum of ∼-0.4 seen in some cells (**Figure 2A**). *d-*scores then rise in female germ cells thereafter, remaining below 0 until E15.5, and only reaching close to parity (i.e. ∼0) at E16.5. Overall the allelic bias observed indicates that a number of genes on the *CAST* X chromosome are clearly silent at E10.5 in female PGCs, which is likely due to the effects of earlier XCI. No allelic expression bias is detected on any of the autosomes at all stages we examined (**Supplementary Figure 2B**). To corroborate these observations further, we chart the expression of *Xist* RNA in our dataset. In keeping with the predictions of XCI, *Xist* is strongly expressed in E11.5 female <u>somatic</u> cells (**Figure 2B**). In comparison *Xist* only expresses at very low levels in E10.5 female PGCs and is downregulated after this stage, consistent with previous studies using RNA FISH (de Napoles et al., 2007; Sangrithi et al., 2017; Sugimoto and Abe, 2007).

**Figure 2.**
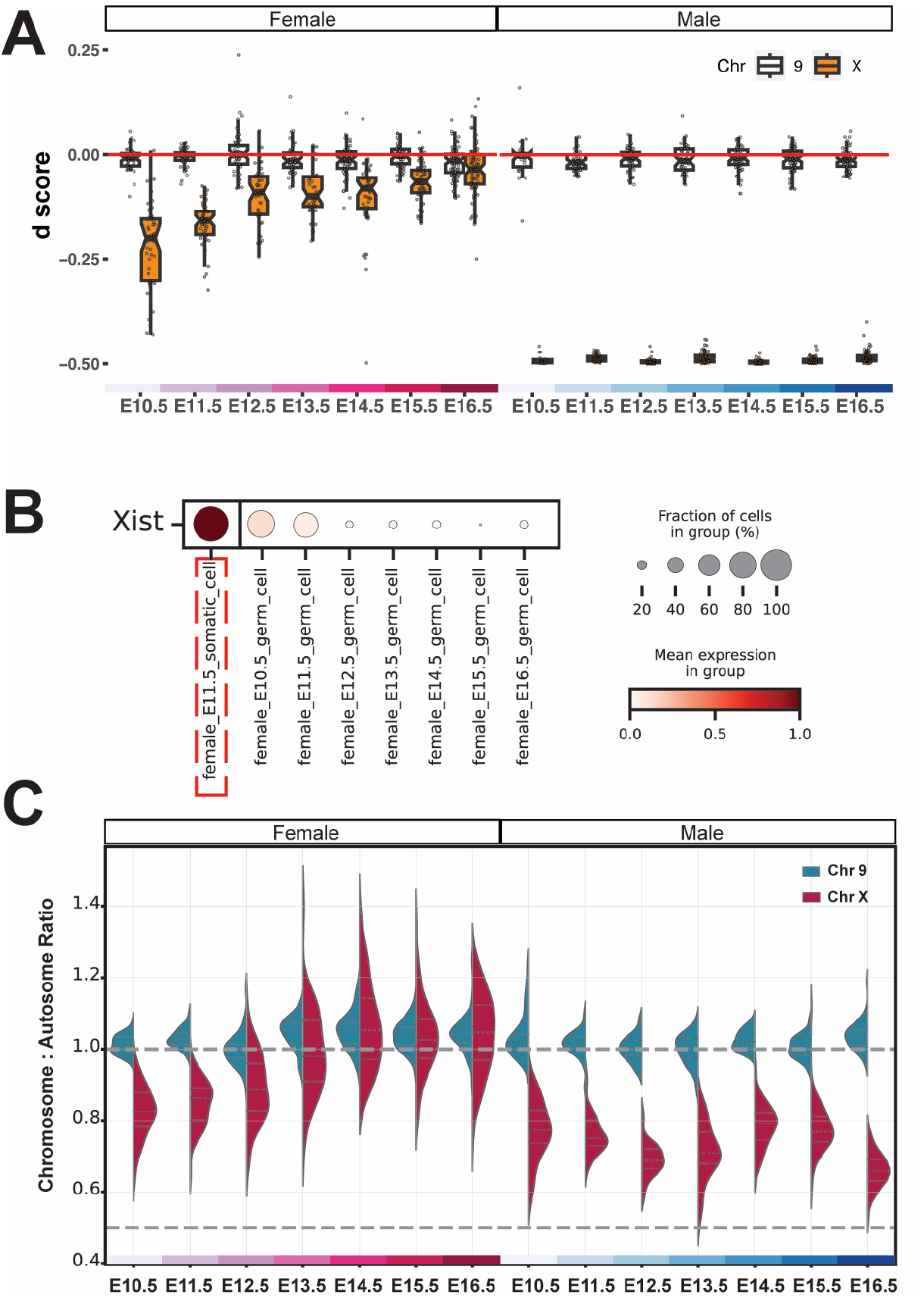
Sexually dimorphic kinetics of X chromosome reactivation and X dosage compensation. A. Boxplot of mean *d*-scores showing allelic balance of chromosome X and 9 during germ cell development in female and male. Each point represents a single cell. B. The dot plot representing *Xist* expression levels in female somatic cells (highted in dashed red square) and in female germ cells at different stages. C. Violin plot representing the ratio of mean read counts of genes on chromosome X (dark red) and 9 (blue) to the mean read counts of all genes across all autosomes for all germ cells at different embryonic stages between female and male.

### Sexually dimorphic dosage compensation states emerge alongside the reactivation kinetics of the X chromosome

We next turned to examining dosage compensation patterns in germ cells. In order to chart global transcriptional output from the X chromosome in relation to the autosomes, we computed X chromosome to autosome expression ratios (X:A) in each cell and plotted these (**Figure 2C)**. We observed that male germ cells consistently have a X:A < 1 over the course of their development. In contrast, female germ cells show increasing X:A ratios, which range above 1 from E13.5 to E16.5. These observations are consistent with previous studies that male germ cells have low X:A ratios (X:A < 1), while females showed an excess of X-gene dosage (X:A > 1) as they enter meiosis (Sangrithi et al., 2017). While X:A ratios in males and females are similar at E10.5, an increase is evident in females thereafter, occurring contemporaneously with XCR as shown earlier (see **Figure 2A**). At E10.5, females have an X:A ratio of 0.82 and males 0.77. X:A ratios show maximal differences between females and males from E14.5 to E16.5 (median values of 1.05 versus 0.66 respectively at E16.5).

### Expression dynamics at individual X gene loci reveal XCR mainly occurs after E12.5

Following stringent filtering we chart the *d-*score for 281 representative X-linked genes subject to XCR, expressing biallelically during female germ cell development (see **Methods**). These are depicted as heatmaps, shown in relation to development age and in context of their genomic location (**Figure 3Ai and ii**). Similarly we also detail their overall expression alongside (**Figure 3Bi and ii**). Together these data depict both allelic balance and expression dynamics for these genes expressing in the female germline from embryonic ages E10.5 to E16.5. From these we surmise that around ∼32% of assayed genes (91/281) express from both alleles (i.e. *d* > -0.4) at E10.5 (**Figure 3C**). This increases to ∼90% of genes expressing biallelically at E16.5 (250/281). Crucially we discover that a significant portion of XCR (∼40%) occurs from E12.5 to E16.5, which is much later than previously appreciated (**Figure 3C**). A number of genes only express later during this time course, with many notably peaking in expression at E16.5.

**Figure 3.**
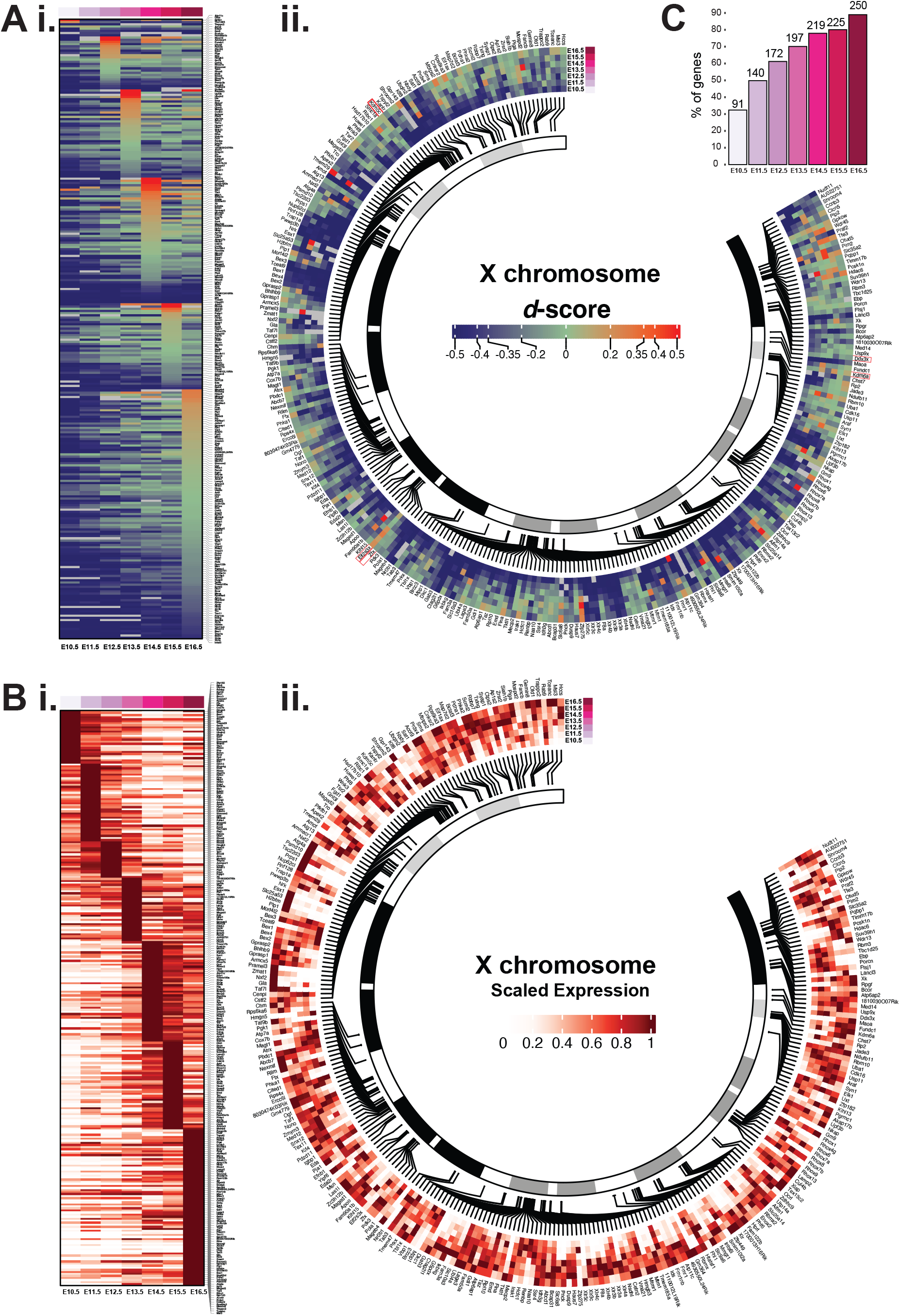
The change in *d*-scores and expression levels of X chromosome linked genes during female germline development from E10.5 to E16.5. A. The heatmap plot of *d*-score showing the allelic balance of 281 X-linked genes; genes are ordered based on (i) reactivation timing according to gestational age and (ii) the genomic location. Five classical XCI ‘escape’ genes (*Ddx3x, Eif2s3x, Kdm5c, Kdm6a* and *Zfx*) are highlighted in a red square. B. The heatmap depicting the expression of 281 X-linked genes; genes are ordered based on (i) reactivation timing according to gestational age and (ii) the genomic location. C. The percentage of X-linked genes expressing biallelically (*d* > -0.4) at each stage.

Further analysis reveals gene expression occurring over the entire length of the X chromosome during germline XCR, even at E10.5. As we observe very low levels of *Xist* expression in E10.5 female PGCs, we examined if XCR could be linked to *Xist* entry sites, by surveying *Xist* RAP data (**Supplementary Figure 3**) (Engreitz et al., 2013). Further to this appraisal we find that neither proximity to *Xist* entry sites nor to the X-inactivation centre (XIC) itself appear to have a significant impact on allelic expression from the reactivating X chromosome in the female germline. This analysis indicates that XCR occurs in a locus-specific manner.

In summary, three important conclusions emerge in this regard. First, we show that the increase in X:A ratios seen during female germ cell development occurs contemporaneously with increasing *d*-scores (i.e. to 0), demonstrating that the increase in expression from the X chromosome is specifically due to biallelic gene expression. Second, allelic balance of the reactivating *CAST* X chromosome and the constitutively active *B6* X reach close to parity at E16.5 when developing oogonia are already in prophase I of meiosis, suggesting that XCR may have an impact in meiosis. Thirdly the protracted reactivation dynamics of the silenced *CAST* X chromosome, beyond *Xist* activity, specifically points toward epigenetic memory from XCI persisting on this chromosome. In all, these results show that germline XCR begins in early PGCs, then proceeds gradually, with most of germline XCR occurring after E11.5 and during female meiosis. While *Xist* RNA expression is notably downregulated after E10.5 in female PGCs, we hypothesise that silencing histone marks deposited as an effect of *Xist* related PRC2 / 1 activity until this developmental age, persists on PGC chromatin (e.g. H3K27me3 and / or H2AK119ub1 deposited consequent to XCI in early PGCs).

### H3K27me3 is enriched on the reactivating X chromosome

To investigate this possibility in greater detail, we next turn to examine how PRC2 dependent H3K27me3 is allelically regulated during germline development. We first optimized low-input <u>c</u>leavage <u>u</u>nder target and release <u>u</u>sing <u>n</u>uclease (CUT&RUN) for H3K27me3 and after validating antibody specificity and reproducibility (**Supplementary Figure 4A and B**), perform low-input CUT&RUN for H3K27me3 on FACS purified germ cell fractions obtained from E13.5 and E16.5 *F*_*1*_ mouse ovaries (Skene and Henikoff, 2017).

H3K27me3 enrichment is seen broadly across the genome, and detected over promoters and gene bodies in female germ cells (**Figure 4A, Supplementary Figure 4C**). Specifically we see a strong enrichment signal over the *Hoxa* gene cluster, which is typically observed in germ cells associated with repression of somatic program (Zheng et al., 2016) (**Supplementary Figure 4D**). From this analysis, we notice a higher enrichment signal for H3K27me3 on the X chromosomes versus the autosomes, more specifically on the *CAST* X chromosome (which is subject to subject to XCI) compared to the *B6* X, clearly more evident at E13.5 versus E16.5 (**Figure 4A**). To confirm differences in H3K27me3 enrichment between the *CAST* and *B6* hemigenomes at each age, we compute a *log2*-ratio of *CAST* over *B6 s*pecific signals in E13.5 and E16.5 germ cells (**Supplementary Figure 4E**). In this manner, we demonstrate a greater enrichment of this mark on the *CAST* X chromosome compared to the *B6* X at E13.5, which we interpret as being due to epigenetic memory of earlier XCI. Crucially, we discover a reduction in the H3K27me3 overall signal in E16.5 oogonia compared to E13.5 oogonia (**Figure 4A**).

**Figure 4.**
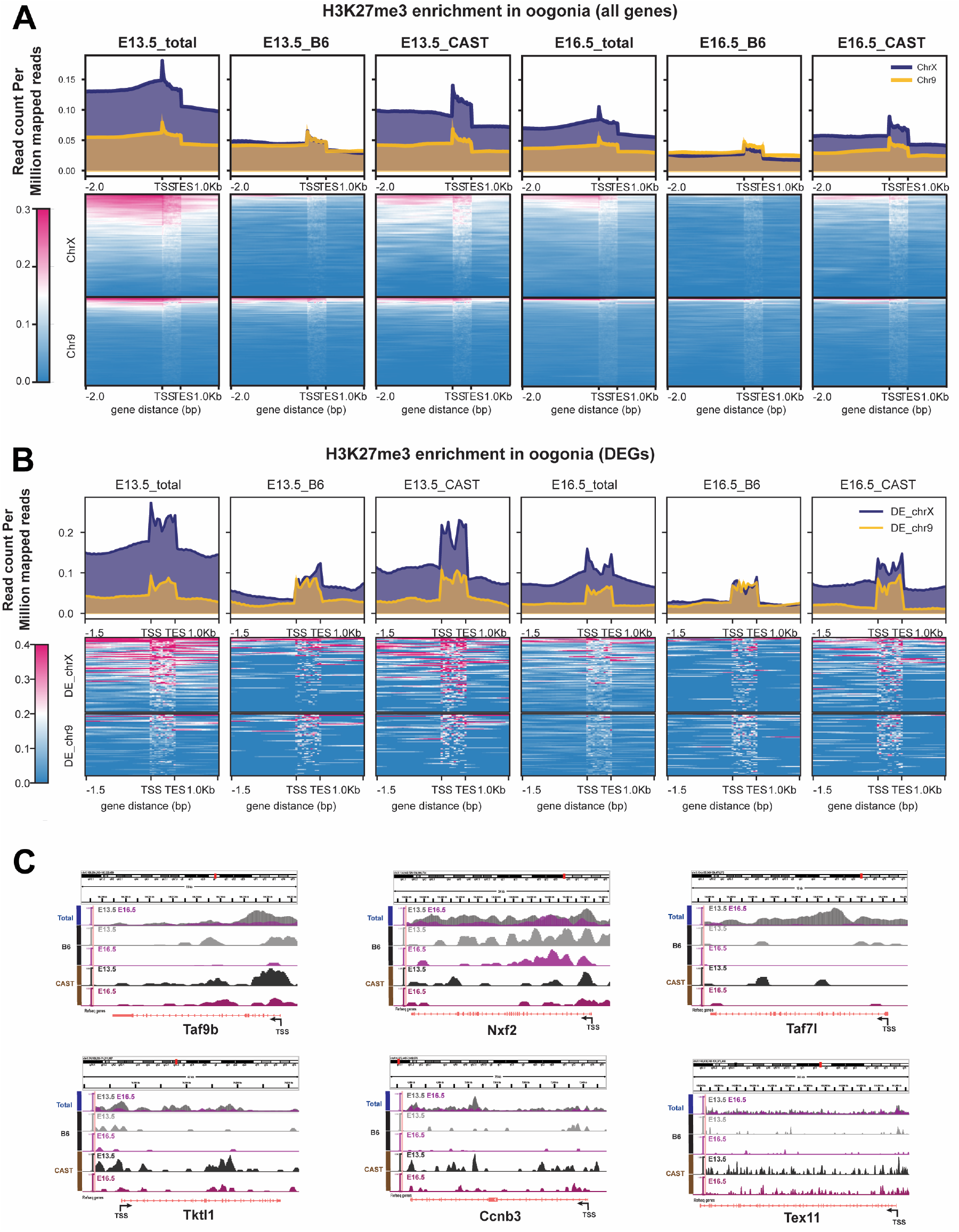
H3K27me3 enrichment remains on the reactivating X chromosome during oogonia development. A. H3K27me3 enrichment signals for all genes on chromosome X and 9 (total), and shown for alleles on the *B6* and *CAST* hemigenomes at E13.5 and E16.5 plotted as heatmaps, with associated summary profiles plotted above. Regions 2kb upstream of transcription start sites (TSSs), gene body and transcription end sites (TESs) are shown. Chromosome X signal is shown coloured orange, and chromosome 9 in blue. B. Genome tracks for H3K27me3 enrichment at E13.5 and E16.5 oogonia for representative genes. C. H3K27me3 enrichment signal at DE genes between E13.5 and E16.5.

Since the reduction in H3K27me3 levels occurs over the same time frame as the increase in X chromosome *d-*score and X:A ratios, we hypothesise that gene activation on the X occurs following removal of this silencing mark. We took two steps to test this directly: 1) we first performed DE analysis to identify genes activated between E13.5 to E16.5 in our RNA-seq dataset (**TableS1**; see **Methods**), and 2) we computed the H3K27me3 signal from CUT&RUN at these DE gene loci in oogonia at E13.5 versus E16.5. In this manner first we identify 76 DE genes on the X chromosome, that increase in expression from E13.5 to E16.5; and 70 DE genes on chromosome 9 were used as autosomal controls. Now returning to CUT&RUN, we examine H3K27me3 levels specifically at these DE genes. Importantly a reduction of H3K27me3 levels is evident at these genes, with the signal change being most pronounced at gene promoters, transcription start sites (TSS) and gene bodies (**Figure 4B**). A number of observations come to light. We detail this reduction at specific DE X-linked gene loci - including *Nxf2, Ccnb3, Tktl1, Taf7l* and *Taf9b*, which have recognized functions in the germ cells and associated with fertility phenotypes (Chotiner et al., 2022; Gura et al., 2020; Pan et al., 2009; Rolland et al., 2011; Wang et al., 2001) (**Figure 4C**).

Overall, we evidence a greater reduction in signal between E13.5 and E16.5 occurring on the X chromosome versus chromosome 9 (**Supplementary Figure 4F**). Looking specifically at the *B6* and *CAST* hemigenomes, we demonstrate a greater reduction in H3K27me3 levels on the reactivating *CAST* X chromosome versus the *B6* X chromosome (or any autosome) at these gene loci (**Figure 4B & Supplementary Figure 4E**,**F)**. These results confirm that expression of these DE genes is specifically associated with the loss of H3K27me3 at promoters, TSS and gene bodies.

### XCR is dependent on Ezh2 activity

Previous studies have shown a vital role for H3K27me3 in regulating sexually dimorphic germline development, regulated by component factors of the PRC2 complex, *Ezh2* and *Eed* (Huang et al., 2021; Lowe et al., 2022). In order to further assess the role of H3K27me3 in this context, we sought to examine data from *Ezh2*-mutant germ cells (**TableS2)** (Huang et al., 2021). From our analysis we notice that a sizeable proportion of genes that are de-repressed in E13.5 *Ezh2*-null oogonia reside on the X chromosome (Huang et al., 2021). Distinctly we see that ∼12.5% of all *Ezh2*-dependent DE genes are X-linked (121 genes with log2folchange > 1; see **Methods**), indicating that *Ezh2* activity significantly regulates gene expression on the X chromosome (**Figure 5A, Supplementary Figure 5B and D**). Again ∼13.2% of DE genes derepressed in E13.5 *Ezh2KO* prospermatogonia were X-linked (48 genes; **Supplementary Figure 5C and D**). These data indicate that Ezh2 dependent H3K27me3 levels on germ cell chromatin regulates the repression of germline genes, orchestrated in a sexually dimorphic manner. We therefore hypothesise that at least some X chromosome genes subject to XCR later (i.e. genes seen to express after E13.5 in our RNA-seq dataset) are directly regulated by H3K27me3 levels. To verify, we first examined the expression of the 121 de-repressed X-linked genes identified in *Ezh2-*null female oogonia, in our scRNA-seq dataset. This analysis indeed confirms these genes are mostly expressed physiologically after E13.5, i.e. from E14.5 to E16.5 (**Figure 5B**). Thus, taken in the context of XCR, these data definitively show that *Ezh2*-dependent H3K27me3 deposition is a functional requirement for the repression of these X-linked genes in E13.5 oogonia, and this silencing mark is then likely reversed for their timely expression observed thereafter. To test this we next turned to directly examine H3K27me3 levels at *Ezh2* regulated genes in oogonia. In keeping with XCR, we observe a greater reduction in H3K27me3 signal at Ezh2-dependent genes on the *CAST* X chromosome identified in our earlier analysis (**Figure 5C)**. In all, our data provide strong evidence that XCR involves the removal of H3K27me3 at specific X-linked gene promoters, TSS and gene bodies by meiosis prophase 1 at E16.5.

**Figure 5.**
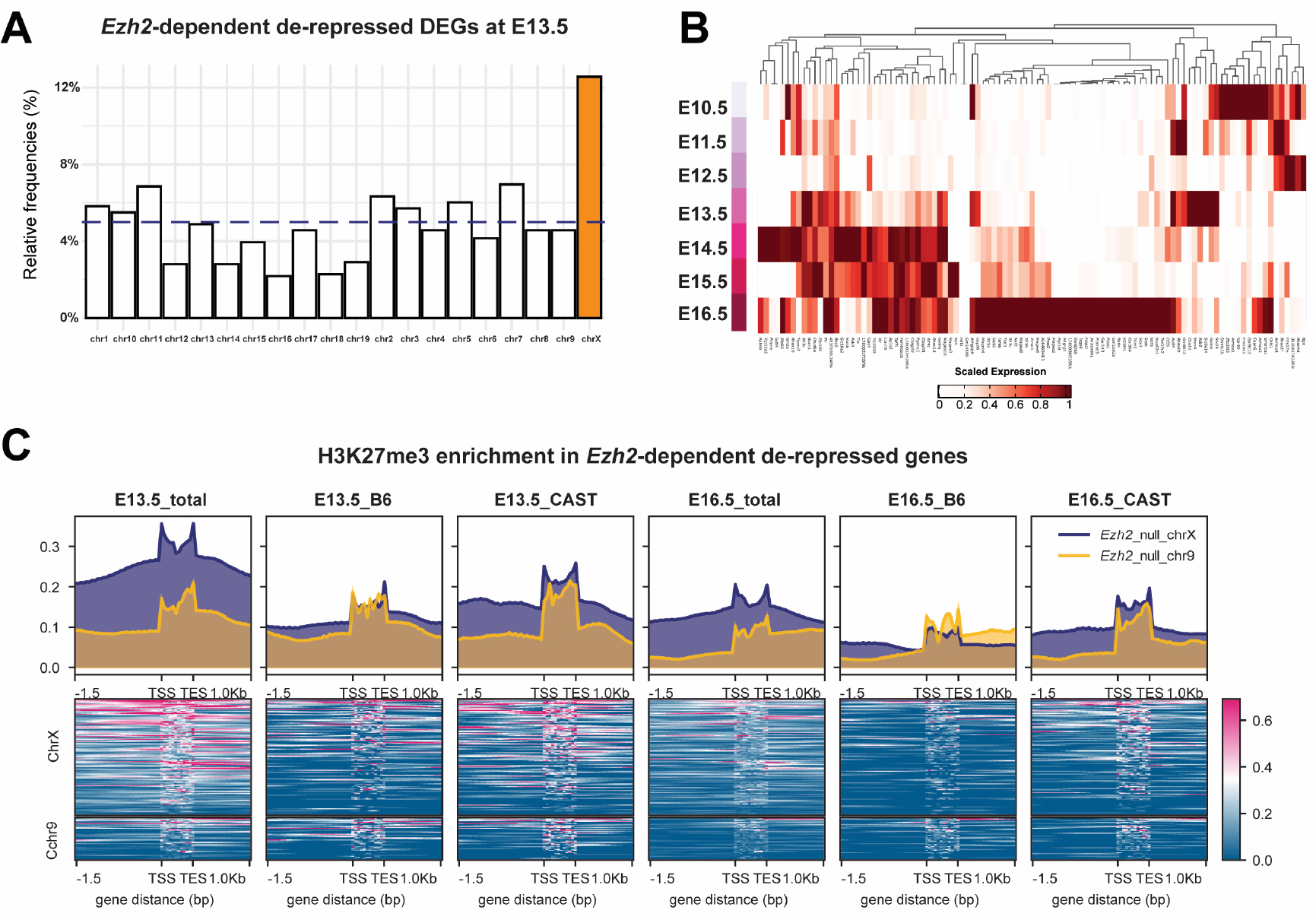
Role of *Ezh2* in X reactivation during oogonia development. (A) The box plot representing the percentages of *Ezh2*-dependent de-repressed genes on each chromosome in the E13.5 oogonia. Chromosome X is shown coloured orange, and gray dotted line indicates the number of expected genes per chromosome if distributed randomly (∼6-8 genes; cf. ∼5-6%). (B) Heatmap showing the expression levels of 121 *Ezh2-*related de-repressed X-linked genes in our scRNA-seq dataset. (C) H3K27me3 enrichment signal for *Ezh2*-dependent genes in our CUT&RUN data, comparing the signal in B6, CAST, and both alleles between E13.5 and E16.5.

## Discussion

In this article we present a highly resolved analysis of X chromosome reactivation kinetics and dynamics using single-cell SMART-Seq2 RNA seq and low-input CUT&RUN at matched time-points. Here our data provides important new insights into gene expression regulation, X chromosome dosage compensation and chromatin dynamics during XCR. While conventional thinking has been that XCR begins early and was mostly complete by the early gonadal phase of germ cells, from our extended work we show that XCR is more protracted than previously appreciated, extends into meiosis, and evidence implicating PRC2 complex (*Ezh2 / Eed / Suz12*) function regulating gene repression on the reactivating X chromosome.

The protracted kinetics of re-establishing transcription on the reactivating X chromosome is striking. We delineate how XCR progresses, and matches with non-canonical X:A dosage compensation patterns observed in the germline. During GWR in female germ cells, X:A ratios exceed 1 from gestational age E13.5. From allele-specific analysis we show that higher X:A ratios in female germ cells are coincident with biallelic gene expression, indicating that X chromosome output arises from the reactivating (*CAST*) X chromosome combined with expression from the constitutively active (*B6*) X chromosome. Our data further highlights a time-window at which near complete reactivation of the X occurs at E16.5 when female germ cells are known to be in late zygotene and early pachytene stage of meiosis.

One of the important findings in this study is the identification of a functional role for H3K27me3 levels in regulating gene expression from the reactivating X chromosome in female germ cells during this developmental time-window. H3K27me3 enrichment on the reactivating X chromosome and its extended presence beyond the expression of *Xist* RNA itself is interesting - notably while *Xist* is downregulated in female germ cells from E10.5, H3K27me3 deposited as a consequence of XCI earlier during PGC development persists, implicating epigenetic memory. This is most likely due to the “self-contained” activity of the PRC2 complex (*Ezh2 / Eed / Suz12*), extending beyond direct *Xist* activity (Yu et al., 2019). There is a precedence for the self-sustaining silencing activity of the PRC complexes on the inactive X (*X*_*i*_), which has also been observed in trophoblast stem cells (Mak et al., 2002; Masui et al., 2023). Here we conclude that XCR does not complete with *Xist* downregulation, and gene silencing on the reactivating X chromosome persists in the absence of *Xist*, due the presence of this repressive histone mark. Self-contained PRC2 activity ensures stable maintenance of H3K27me3 mediated gene silencing, mitigating against cell-cycle dependent dilution of this mark. Female germ cells cycle mitotically until ∼E13.5 at which point they commit to meiosis. In the absence of cell-cycle dependent dilutional effects beyond this stage in the female germ line, the loss of H3K27me3 signal observed thus points to the existence of active processes to reverse H3K27me3 silencing at these loci. In this regard, future studies could be aimed at identifying and evaluating such mechanisms involved in reversing H3K27me3 mediated gene silencing in the germline.

The presence of H3K27me3 on the reactivating X chromosome crucially also accounts for the dynamic X:A dosage compensation pattern seen in XX germ cells. The relative overexpression of X-genes seen from E13.5 in females, observed by X:A ratios ranging above 1, could be attenuated by this silencing mark (cf. related observations made in the male by Lowe et al. (2022)).

It is unexpected that the reactivating X chromosome remains enriched for H3K27me3 over the constitutively active X in female germ cells in the context of meiosis. PRC2 is known to act as a factor to maintain repression of distinct gene types: e.g. developmentally regulated genes, protein-coding genes that are repressed (including germ line genes) and imprinted genes. Thus, the control of PRC2 activity may be developmentally regulated to promote meiosis via the timely expression of germline genes in a sex-specific manner. Three further points also come up in this regard. First the asymmetric distribution of H3K27me3 on the X chromosome at the point of meiotic entry at E13.5 merits further investigation (i.e. in chromosome pairing, synapsis and segregation). Second the regulation of heterochromatin more generally during female meiosis would also be of interest as it highlights possible roles for other silencing marks on the inactive X (e.g. H2AK119ub1 and H3K9me3). And third it of importance to understand the mechanisms involved in the reversal of silencing on the X, to achieve expression. Furthermore, from an evolutionary perspective it would be of interest to compare XCR in other mammals such as humans to understand conservation and / or divergence in these fundamental epigenetic processes.

Despite the well-known complex dynamics of XCR, notably important differences are apparent with our *in vivo* appraisal of XCR and previous studies using *in vitro* derived PGC models of XCR, where incomplete XCI had occurred in PGCLCs further limiting meiotic potential (Severino et al., 2022). This was confirmed in a recent manuscript using a random XCI model in early *in vivo* PGCs (Roidor *et al*. 2023), where PGCs had mostly maintained an inactive X chromosome at E9.5, with a limited number of genes undergoing XCR thereafter. From the experimental strategy employed in this study using *F*_*1*_ female (X_*CAST*_X_*B6-XistΔ*_) embryos, gestational ages covered, we are able to demonstrate many more genes subject to XCR. This supports the finding that a number of genes are orchestrated to undergo XCR with important functional roles therein in development of the female germ line.

In conclusion we present new evidence that challenge existing paradigms in X chromosome biology, showing that XCR extends much later than previously appreciated in female germ cells. Our work further connects PRC2 activity beyond a sex specific role in the germline, more directly in regulating X chromosome activity in promoting female meiosis (Huang et al., 2021; Lowe et al., 2022).

## Supplementary Figure legend

**Supplementary Figure 1. Quality control of scRNA-seq data**

(A) Dot plot of total counts, number of genes expressed, and the percentage of mitochondrial reads in the scRNA-seq data in this study. Each dot represents a single cell.

(B) Summary of cells represented in scRNA-seq dataset.

(C) The expression level of Y-chromosome encoded genes across different embryonic stages between male and female.

(D) The expression level of key genes across cells, visualised on the *t-SNE*.

**Supplementary Figure 2. The allelic expression of chromosomes in female cells (relates to Figure 2)**

(A) The boxplot of *d*-scores of chromosome X and 9 in E11.5 female germ cells and somatic cell.

(B) The boxplot of *d*-scores of all autosomes and chromosome X in oogonia from E10.5 to E16.5. Each dot represents a single cell sample.

**Supplementary Figure 3 (relates to Figure 3)**

The heatmap plot of *d*-scores for of 281 X-linked genes; genes are depicted based on the genomic location. The *Xist* RAP-seq signal is represented in blue in the figure, and putative *Xist* entry sites as red dots. *Xist* / XIC location is marked as shown (high signal for *Xist* over the XIC is not shown).

**Supplementary Figure 4. CUT&RUN results (relates to Figure 4)**

(A) The H3K27me3 antibody verification: heatmap showing the H3K27me3 antibody specificity validated using SNAP-CUTANA K-MetStat Panel. Representative immunofluorescence image of α-H3K27me3 (magenta), α-Ddx4 (cyan) and endogenous GFP in an E13.5 ovary. White arrows indicate Barr bodies in somatic cells.

(B) The heatmap representing the spearman correlation of reads counts generated from CUT&RUN samples.

(C) The genome tracks showing H3K27me3 enrichment signal over *Hoxa* gene cluster in E13.5 and E16.5 oogonia.

(D) The distribution of H3K27me3 peaks across functional annotations, including promoter, exons, introns, downstream, and distal intergenic regions in E13.5 and E16.5 germ cells.

(E) The box plots showing the *log2*-ratio of H3K27me3 enrichment signal in *CAST* to *B6* alleles in all the chromosomes. Chromosome X is highlighted in orange colour and 9 in blue (relates to **Figure 4A**).

(F) The *log2*-ratio of H3K27me3 enrichment depicting change in signal in E13.5 over E16.5 germ cells at DE genes. Plots show higher enrichment at E13.5, especially at CAST specific alleles (relates to **Figure 4C**).

**Supplementary Figure 5. Analysis of *Ezh2*-null germ cells scRNA-seq (relates to Figure 5)**

(A) Principal component analysis (PCA) of RNA-seq data from *Ezh2*-null (conditional knockout) and control male and female E13.5 gem cells.

(B) Volcano plot showing the differentially expressed genes (adjusted p-value <0.05, log2FC>1 or log2FC<-1, orange dots) between *Ezh2*-null and control female germ cells. The X-linked DEGs were highlighted in dark blue (n=121).

(C) Volcano plot showing the differentially expressed genes (adjusted P value <0.05, log2FC>1 or log2FC<-1, orange dots) between *Ezh2*-null and control male germ cells. The X-linked DEGs were highlighted in dark blue (n=48).

(D) Summary of DE genes between *Ezh2*-null and Control mice.

## Acknowledgements

We thank Dr. James Turner (Francis Crick Institute) for enabling pilot experiments / sharing relevant mouse lines, helpful discussions, critical reading of the manuscript. We thank members of the Sangrithi lab for discussions and helpful feedback. We thank staff at the KCL Biological Services Unit for assistance with animal husbandry and welfare. Access to high-performance computing resource for the Sangrithi lab is obtained from King’s College London, King’s Computational Research, Engineering and Technology Environment (CREATE). This work was supported by the Francis Crick Institute which receives its core funding from Cancer Research UK (CC2052), the UK Medical Research Council (CC2052), and the Wellcome Trust (CC2052). MS is supported by the Wellcome Trust (222052/Z/20/Z).

## Contributions

YL designed and performed all the experiments and maintained the mouse lines. YL and XZ collected the samples. Bioinformatic analysis were performed by PM and supervised by MS. CASS performed analysis of *Ezh2*-null data and immunofluorescence microscopy. DS performed FACS. MS designed and supervised all experiments, wrote the manuscript, maintained all university compliances, and attained funding for the experiments.

## Data availability

Single-cell RNA-seq data and CUT&RUN data will be publicly available as of the date of publication in a peer-reviewed journal.

## METHOD DETAILS

### Biological Samples and the Ethical Use of Animals

Use of experimental mice was undertaken in strict accordance with the Animals (Scientific Procedures) Act 1986 subject to local ethical review and carried out under UK Home Office license. Mice were maintained on a 12:12 hour light : dark cycle at 22 ± 2 °C, with food and water available *ad libitum. Oct4-EGFP* mice are maintained on a C57B6 background and used to mark / isolate fluorescently marked germ cells (Yeom et al., 1996; Yoshimizu et al., 1999). *XistΔ* mice are also maintained on a C57B6 background (Marahrens et al., 1997) and maintained with the *Oct4-EGFP* line. *Mus castaneus* (*CAST / EiJ*; RRID*:IMSR_JAX:000928*) were purchased from The Jackson Laboratory (JAX).

Experimental *F*_*1*_ embryos were generated by crossing *Xist*^*+/Δ*^ (B6) *dams* also carrying *Oct4-EGFP* transgene, and *CAST sires*, producing *F*_*1*_ female (XX^XistΔ^) and males (XY) that were analyzed. Typically timed natural matings were set up at around 17:00 hrs, by placing female mice into with the male. Females were checked for the presence of a vaginal plug indicating a mating has occurred, with noon on the day the vaginal plug taken as E0.5 and was separated from the male.

#### Genotyping

PCR genotyping was performed on extracted DNA. The primer information is presented in **TableS3**.

##### Sample collection and processing

*F*_*1*_ embryos were harvested daily at E10.5 to E16.5 from the date of the vaginal plug, with further morphological verification at dissection. The embryonic urogenital complexes were carefully removed under the stereomicroscopy, and washed in cold DPBS twice. For transcriptomic and CUT&RUN, the embryonic gonads were separated from adjacent mesonephroi and placed into 1.5 mL Eppendorf tube with 300uL DPBS.

#### FACS Isolation of germ cells

To obtain a single-cell suspension for FACS, isolated gonads were digested in the HBSS buffer containing Collagenase Type 1 (200U/mL), Dispase II (2.4 U/mL), and CaCl_2_ (1mM) at 37°C. Tubes were shaken every 2-3 mins until the tissue fully dispersed. Enzymatic digestion was then neutralisation with DMEM, containing 10% foetal bovine serum, followed by centrifugation at 1000g, at 4°C for 4 minutes. The cell pellet was then resuspended in cold FACS buffer (1% BSA in HBSS, with 25 mM HEPES, 1mM EDTA, and PI buffer) and filtered through a 40μM strainer. EGFP fluorescence was used to isolate the fluorescently marked germ cells through FACS. Live cells germ cells (PI negative, GFP positive) and somatic cells (PI negative, GFP negative) were sorted by florescence-activated cell sorting (FACS) using ARIA III flow cytometer (BD Bioscience).

For single cell RNA-seq, individual cells were sorted into 96-wells plates containing lysis buffer. Plates were sealed and centrifuged immediately (700g, 10 seconds) following the sort and stored at -80°C until needed. For CUT&RUN experiments, bulk sorting was performed to collect purified germ cell and / or somatic cell fractions separately into 300μL of collection buffer (5% BSA-HBSS) and used fresh.

##### SMART-Seq2 library preparation and sequencing

SMART-seq2 method was applied to generate the full-length cDNAs. Briefly, a single cell (∼0.3μL) was sorted into the 96 -wells plates containing 4 μL of lysis buffer (a ribonuclease (RNase) inhibitor, dNTPs, and oligo-dT oligonucleotides). Cell lysis was performed by incubating samples at 72 °C for 3 mins and put back on ice immediately. Then, reverse transcription (RT) reaction was carried out by adding 0.5μL Maxima H-reverse transcriptase (200 U / μL), 0.25μL RNA inhibitor (40 U / μl), 2μL Maxima-H RT buffer (200U / μL), 2μL Betaine (5M), 0.06μL MgCl_2_ (1M), 0.1μL TSO (100 μM), and nuclease-free water to make a total of 10μl reaction. The reaction program of Maxima H reverse transcriptase was 90 mins at 50°C, followed by 5 mins enzyme activation at 85°C; the reaction was then hold at 4°C. The pre-amplification was then performed by adding 12.5μL KAPA HiFi HotStart, and 2.5μL nuclease-free water. 20 PCR cycles were used, and the PCR cycle was set as follow: 98°C for 3 mins, 20 cycles of 20s at 98°C, 15s at 67°C, and 6 mins at 72°C. The final elongation was performed at 5 mins at 72°C. After purification with AMPure XP beads (Beckman Coulter) (in a ratio of 0.7:1), the PCR product was quality checked (QC). Tagmentation of 0.3ng cDNA was carried out by using the Illumina Nextera XT DNA sample preparation kit in a reaction volume of 4 μL (2μL of tagmentation DNA buffer (TD, 2 x); 1μL of Amplicon tagmentation mix, and 1μl of diluted cDNA), followed by stripping Tn5 transposase off with 1μL of NT buffer added. Finally, amplification of adapter-ligated fragments was performed by adding 3μL Nextera PCR master mix, and 2μL of Index primer combination. After purification with AMPure XP beads (in the ratio of 0.6:1), the final cDNA library was quantified and QC-ed.

The concentration of PCR product was measured by using Qubit dsDNA assay kit (Invitrogen), and size distribution was checked using an Agilent high-sensitivity chip (D1000) on 4200 TapeStation System. Overall, 741 scRNA-seq libraries were prepared and 2x150 bp paired-end (PE) reads was performed on an Illumina Novaseq 6000 instrument. Where possible we aimed to obtain and sequence a minimum of 24 cells from each embryo.

### RNA-seq data analysis and processing

All processing and analyses were performed on the high-performance computing (HPC) platform King’s Computational Research, Engineering and Technology Environment ([CREATE](https://doi.org/10.18742/rnvf-m076)). Fastq files containing reads from single-cell RNA seq libraries were subject to quality control ([fastqc](https://www.bioinformatics.babraham.ac.uk/projects/fastqc/)), and trimmed for adapter content using [Trimgalore](https://github.com/FelixKrueger/TrimGalore). Using the SNPsplit tool, SNPs positions for *C57B6NJ* and *CAST / EiJ* in the GRCm38.p6 reference mouse genome ([Ensembl](https://nov2020.archive.ensembl.org/Mus_musculus/Info/Index)), were first N-masked to eliminate mapping-bias between alleles (Krueger and Andrews, 2016). An N-masked genome index was then generated using *STAR* (Dobin et al., 2013; van de Geijn et al., 2015). Trimmed fastq files were subsequently aligned to the N-masked reference using the *STAR* aligner.

Total gene-level expression counts were then generated using *featureCounts* (*Subread*) and used in downstream analyses (Liao et al., 2014). Similarly allele-specific counts for alignments specific to *B6* and *CAST* were generated using split bam files generated by SNPsplit.

Total counts were imported into *SCANPY* for further specific analyses including normalization, dimensionality reduction and statistical approaches (Wolf et al., 2018). Cells kept fulfilled the following criteria - min_genes > 3500, max_genes < 10000, min_counts > 500000, mitochondrial counts < 5%. Genes kept for analysis fulfilled the following criteria - min_cells = 5. Following this, 681 (out of 738) high-quality single-cells transcriptomes were retained for further processing with each time point represented by a minimum of 20 cells derived from each embryo (please see **Supplementary Table 1B**). *In toto* 681 cells and 21174 expressed genes were analysed.

### Allele-specific analyses

Custom scripts in R were used to derive allelic-ratios and *d-scores* (*d*) from count matrices output from *featureCounts. CAST* allelic ratios were calculated as *CAST* / (*B6* + *CAST*), and *d* score *= CAST / (B6 + CAST) - 0.5)* was calculated for each gene and / or each cell. Reads to the *CAST* X chromosome in male cells likely arising from technical artifacts of singular mis-annotated SNPs, were not considered. For XCR analysis, genes with counts detected in at least 2 or more cells, and 5 fragment counts or more at each gene locus were kept for XCR analyses.

#### Cleavage Under Targets and Release Using Nuclease (CUT&RUN)

For each CUT&RUN reaction, typically ∼5000 cells were used, pooled from different embryos of identical genotype in the same litter. Here we applied a CUT&RUN method modified from previous protocol (Derek Janssens, Steven Henikoff 2019. CUT&RUN: Targeted in situ genome-wide profiling with high efficiency for low cell numbers. protocols.io https://dx.doi.org/10.17504/protocols.io.zcpf2vn). Sorted cells were washed with Wash Buffer (20 mM HEPES, pH7.5, 150 mM NaCl, 0.5 mM spermidine and a Roche complete tablet per 50 ml), and then bound to activated Concanavalin A (ConA) -coated magnetic beads. After 10 mins of binding, cells were permeabilized and incubated in 50μL Antibody Buffer (Wash buffer containing 0.01% Digitonin and 0.5μL of specific primary antibody) at 4°C overnight on a nutator. For the negative control, anti-Rabbit IgG was added instead of primary antibody. Next day, after 2 washes with Digitonin buffer (Wash Buffer containing 0.01% Digitonin), beads were resuspended in pAG / MNase buffer (50μL Digitonin buffer with 2.5μL of pAG / MNase reagent (EpiCypher) and incubated at room temperature (10 mins) on a nutator for pAG-Mase binding. After washing away pAG / MNase with Digitonin buffer, the chromatin digestion was performed by adding 1μl of 100mM CaCl_2_ to each reaction and incubating at 4 °C for 2 hours on a nutator. Digestion was then stopped with the addition of 33μL Stop Buffer (340 mM NaCl, 20mM EDTA, 4mM EGTA, 50μg / mL RNase A, and 50μg / mL glycogen) and the cleaved chromatin was released to supernatant by incubating at 37°C for 10 mins. DNA was then purified using a DNA clean-up column (Monarch® PCR & DNA) following the manufacturer’s instructions, and used for library preparation (NEBNext^®^ Ultra™ II DNA Library Prep Kit for Illumina^®^). Specifically, the concentration of adaptor (reduced to 3μM) and the cycle number of PCR amplification (15 cycles) were optimised in our experiment considering of the low input of DNA. Finally, the clean-up of PCR product was carried out by using AMPure X beads (Beckman Coulter) (in a ratio of 0.9:1). After measuring the quantity of library and checking the size distribution, libraries were pooled and sequenced to produce 2 X 50bp PE sequencing on Illumina NextSeq 2000 instrument.

We determined the specificity for the antibodies used, using the SNAP-CUTANA K-MetStat Panel (EpiCypher, US). 2μL of 1:20 diluted solution was applied to ConA-immobilized cells prior to the addition of antibody. As per the manufacturer’s instructions, reads were matched to the unique DNA barcodes in the panel and normalized to either on-target or total counts. Antibody enrichment of the expected spike-in nucleosome target confirmed the integrity of the H3K27me3 antibody used in our experiment, while the IgG control antibody showed no preferential enrichment for any nucleosome as expected (**supplementary Figure 4A**). In addition, we validated this antibody by immunostaining of H3K27me3 on E13.5 embryo gonads (**supplementary Figure 4A**).

##### CUT&RUN data processing

In short, *fastq* files were pre-processed for quality and trimmed (min length >35). Reads were aligned to the N-masked reference genome using *STAR* with spliced-alignments turned off. Subsequent steps in the analyses were performed using the *deepTools* package (v3.5.0) (Ramirez et al., 2016). Pearson correlations were computed and plotted to visualize cross-replicate validity (*multiBamSummary*), before proceeding to merge bam files at each time-point. Reads across the genome were counted in 10kb-size bins for each library, and effective library sizes were calculated using *csaw*, with and the TMM method applied to compute normalization factors (Lun and Smyth, 2014; Robinson et al., 2010). *Bigwig* files were generated from the *bam* files using *bamCoverage* with settings “-binSize 100 --smoothLength 1000”. Signals were further computed / visualized using deepTools’ functionality (e.g. *computeMatrix, plotHeatmap*), and to plot enrichment profiles relating to each timepoint.

##### Quantification and statistical analysis

Statistical analyses were performed in R statistical computing playform (https://www.r-project.org/) or Python. Statistical tests and the cut-off values used for each analysis are described in each methods subsection. p<0.05 was considered significant for all tests. For the biological replicated, in the scRNA-seq experiment, at least 2 embryos and 24 scRNA-seq libraries / embryo were generated at each time-point. In the CUT&RUN experiment, 2 biological repeats were carried out at each time-point.

