## Supplementary_Figures for "Single cell RNA-seq reveals protracted germ line X chromosome reactivation dynamics directed by a PRC2 dependent mechanism"

A

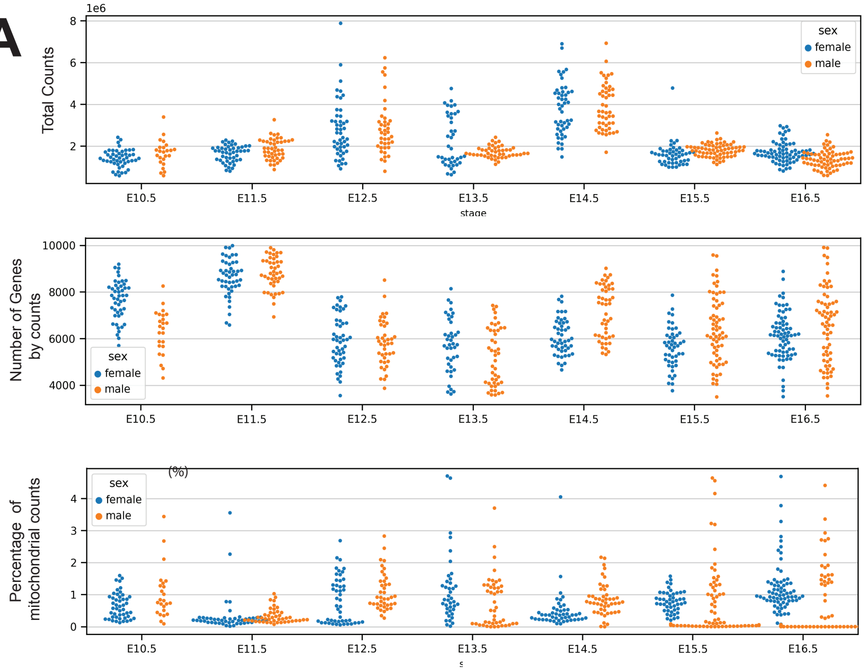


B

| Cell type | Embryo stage | Sex | Embryo number | scRNA libraries | scRNA library past QC |
|---|---|---|---|---|---|
| Germ cell | E10.5 | Female | 2 | 48 | 47 |
| | | Male | 1 | 48 | 24 |
| | E11.5 | Female | 2 | 48 | 47 |
| | | Male | 2 | 48 | 44 |
| | E12.5 | Female | 2 | 48 | 48 |
| | | Male | 2 | 45 | 40 |
| | E13.5 | Female | 2 | 48 | 40 |
| | | Male | 2 | 48 | 45 |
| | E14.5 | Female | 2 | 48 | 48 |
| | | Male | 2 | 48 | 48 |
| | E15.5 | Female | 1 | 48 | 47 |
| | | Male | 3 | 72 | 65 |
| | E16.5 | Female | 2 | 72 | 71 |
| | | Male | 3 | 72 | 67 |
| | Total | | 28 | 741 | 681 |
| Somatic cell | E11.5 | Female | 2 | 24 | 24 |
| | | Male | 2 | 24 | 24 |

C

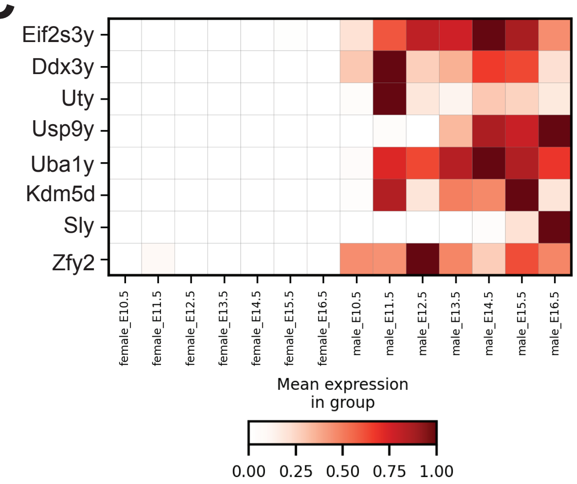


D

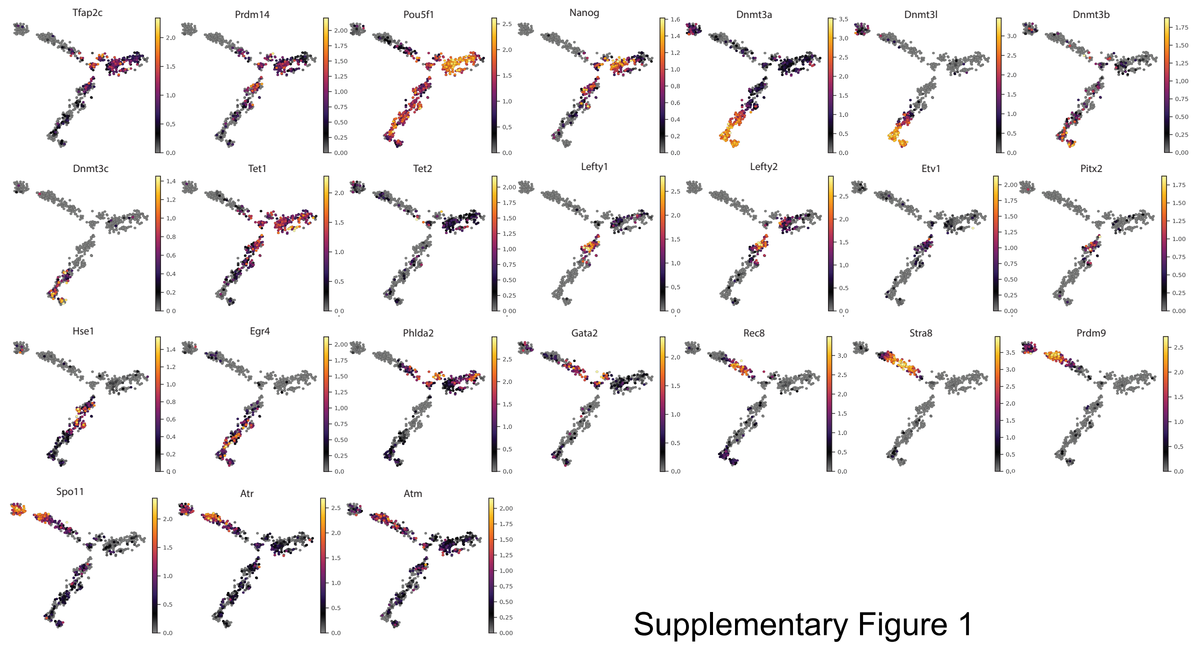


Supplementary Figure 1

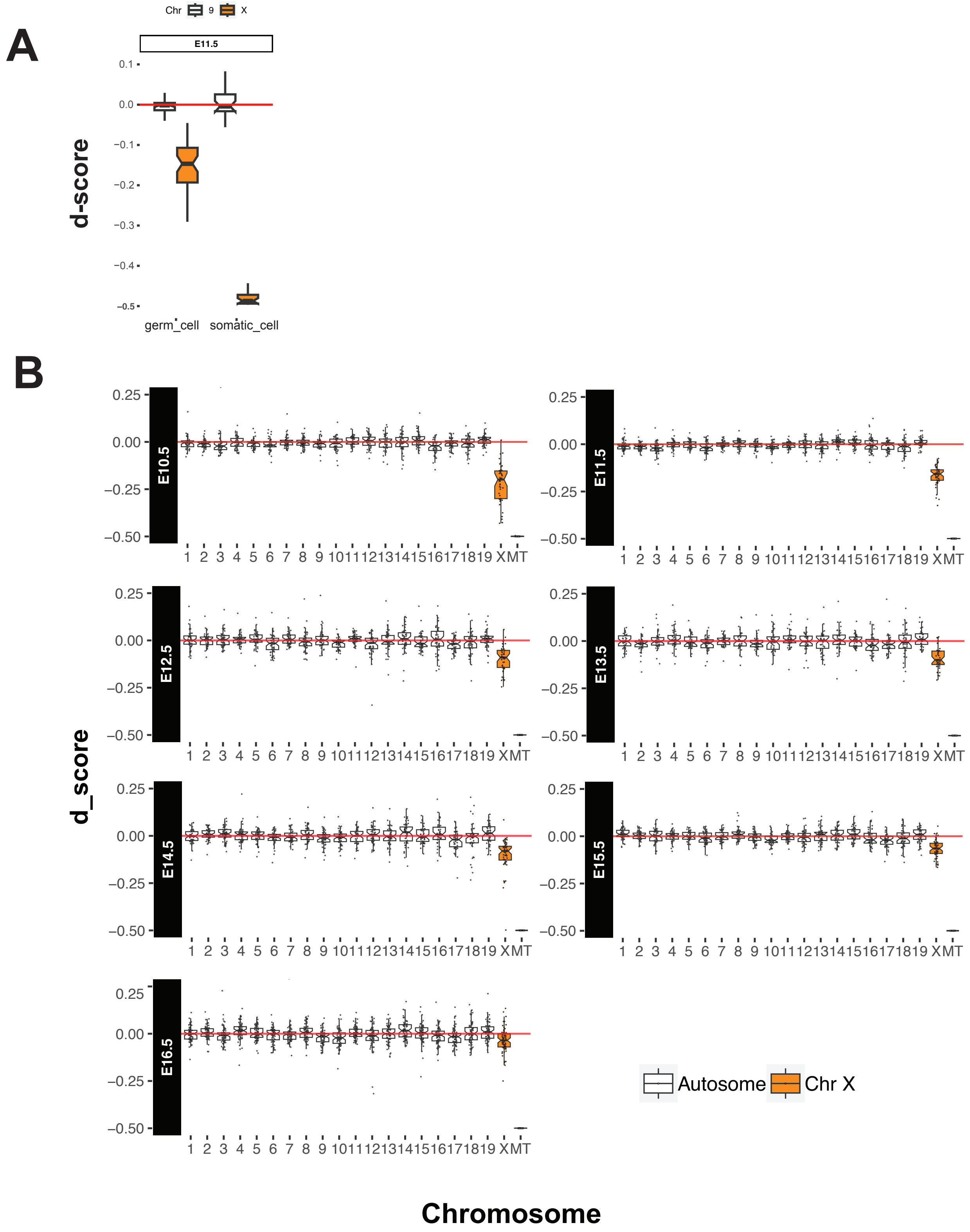


Supplementary Figure 2

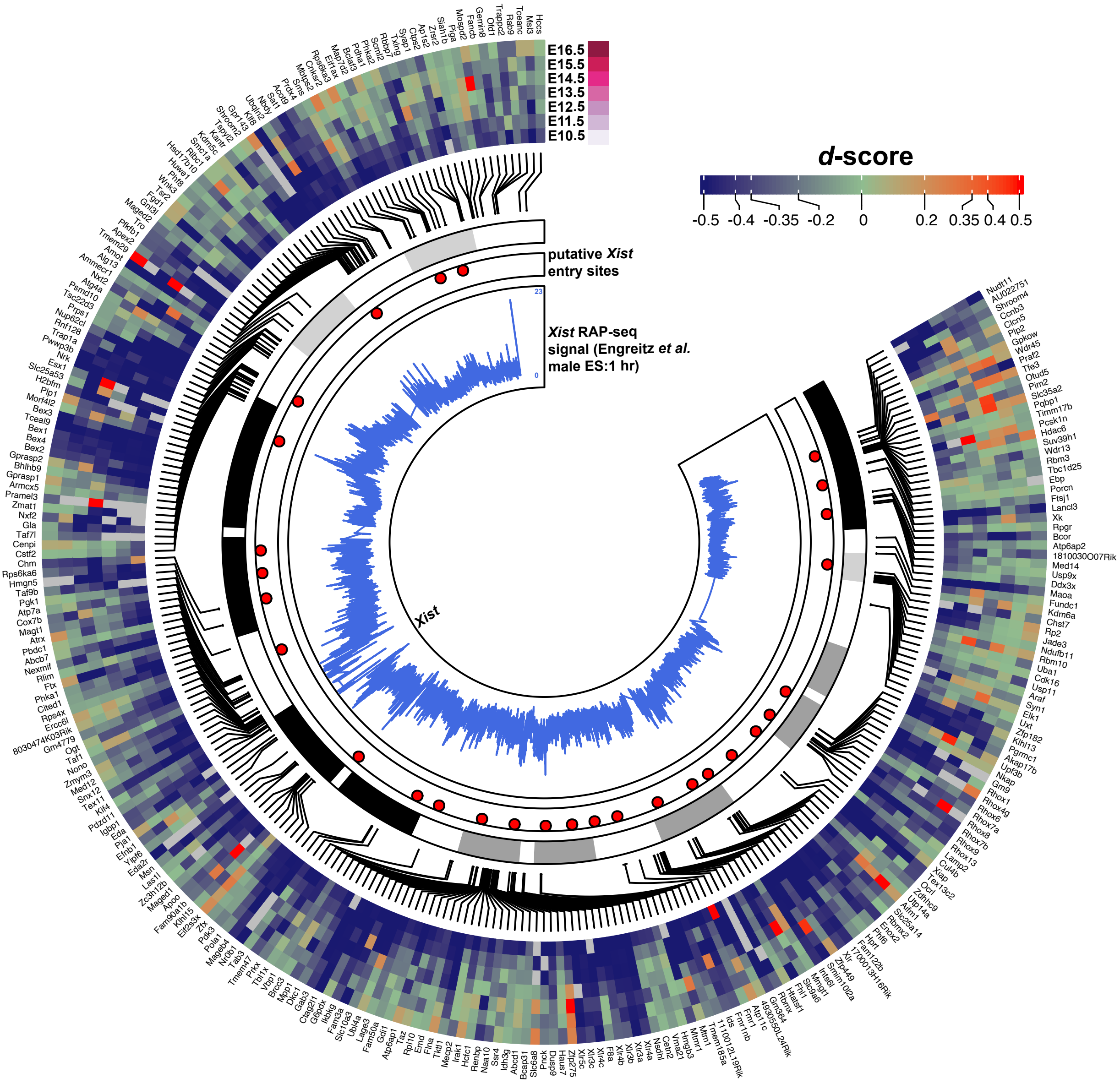


Supplementary Figure 3

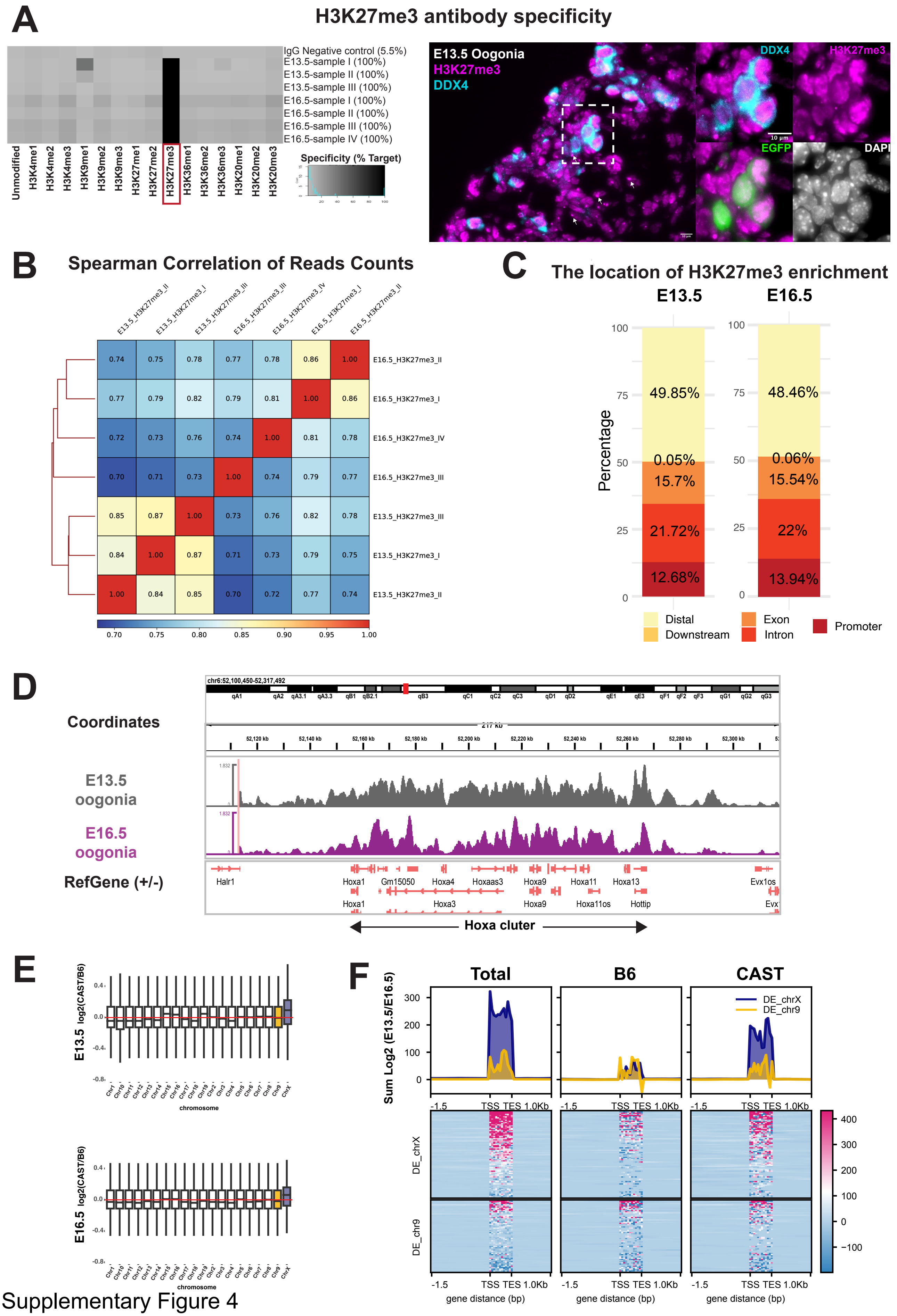


Supplementary Figure 4

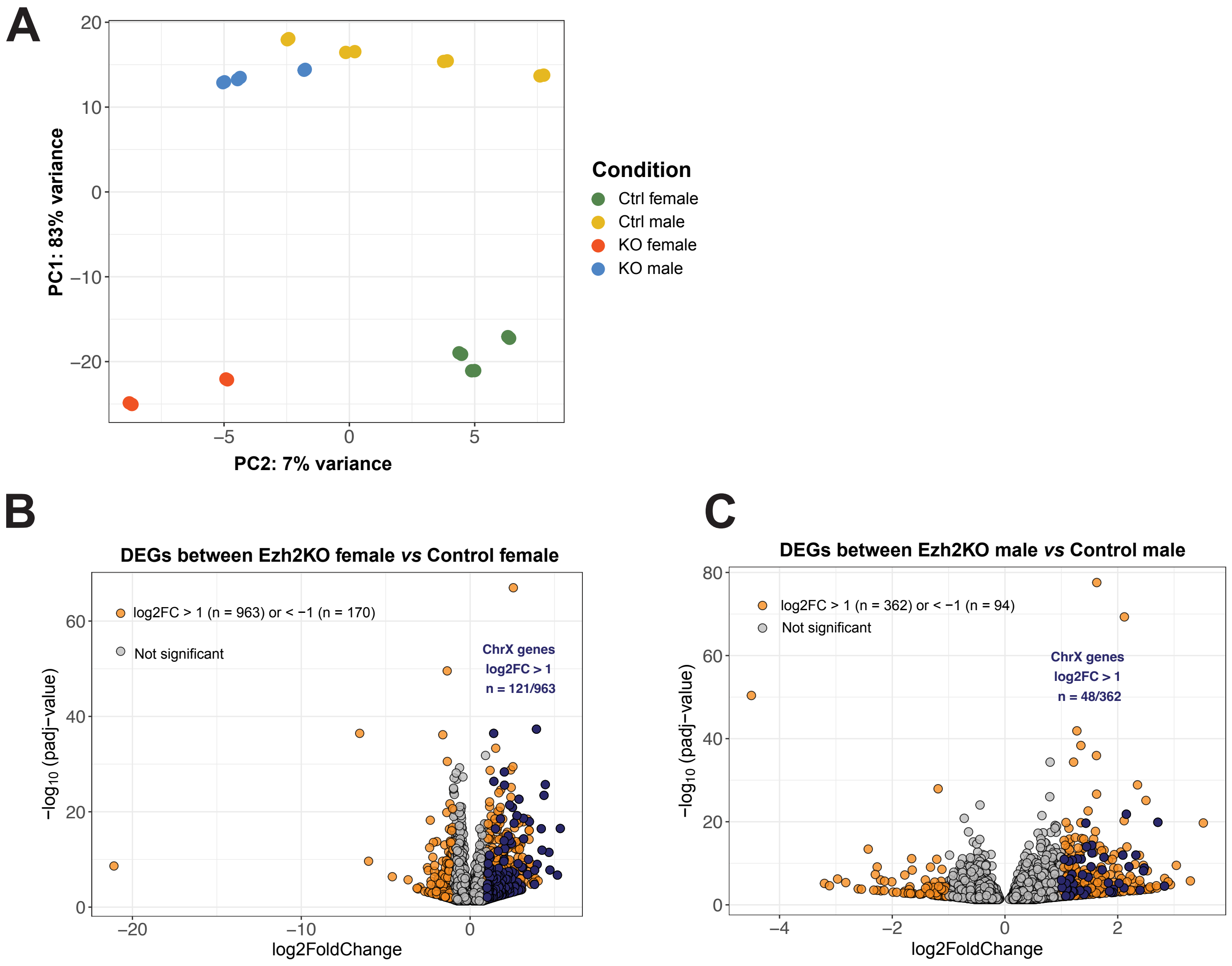


**D**

| Criteria for DE genes | Male Ezh2KO vs Ctrl | Female Ezh2KO vs Ctrl |
|---|---|---|
| padj < 0.05 | 3980 | 5545 |
| padj < 0.05, log2FoldChange > 1 | 362 | 963 |
| padj < 0.05, log2FoldChange < -1 | 94 | 170 |
| padj < 0.05, log2FoldChange > 1 on the ChrX | 48 | 121 |
| padj < 0.05, log2FoldChange < -1 on the ChrX | 3 | 0 |

Supplementary Figure 5
